# Gamma-band resonance of visual cortex to optogenetic stimulation

**DOI:** 10.1101/135467

**Authors:** Jianguang Ni (倪剑光), Christopher Murphy Lewis, Thomas Wunderle, Patrick Jendritza, Ilka Diester, Pascal Fries

**Affiliations:** Ernst Strüngmann Institute (ESI) for Neuroscience in Cooperation with Max Planck Society, Deutschordenstraβe 46, 60528 Frankfurt, Germany; International Max Planck Research School for Neural Circuits, Max-von-Laue-Straβe 4, 60438 Frankfurt, Germany; Donders Institute for Brain, Cognition and Behaviour, Radboud University Nijmegen, Kapittelweg 29, 6525 EN Nijmegen, Netherlands

## Abstract

Activated visual cortex typically engages in neuronal synchronization in the gamma-frequency band (30-90 Hz). Gamma-band synchronization is related to cognitive functioning, and its mechanisms have been extensively investigated, predominantly through in-vitro studies. To further elucidate its mechanisms in-vivo, we performed simultaneous optogenetic stimulation and electrophysiological recordings of visual cortical areas 17 and 21a in the anesthetized cat. Viral transfection with AAV1 or AAV9 under a CamKIIα promoter led to robust Channelrhodopsin-2 (ChR2) expression. Immunohistochemical analysis showed that all ChR2-expressing neurons were negative for Parvalbumin, consistent with predominant or exclusive expression in excitatory neurons. Optogenetic stimulation used primarily surface illumination directly above the transfected and recorded cells. Stimulation with constant light led to strong and sustained gamma-band synchronization with strength and bandwidth similar to visually induced gamma. Rhythmic stimulation with light-pulse trains or sinusoidal light modulation revealed strongest resonance for gamma-band frequencies. Gamma resonance was confirmed by optogenetic white-noise stimulation. White-noise stimulation allowed the quantification of the transfer function between the optogenetic stimulation and the local field potential response. This transfer function showed a dominant peak in the gamma band. Thus, we find that visual cortical circuits resonate most strongly to gamma-band components in their input. This resonance renders both the sensitivity to input, and the output of these circuits, selectively tuned to gamma.

**Significance Statement:** Activated groups of cortical neurons often display rhythmic synchronization in the gamma-frequency band (30-90 Hz). Gamma-band synchronization is particularly well studied in visual cortex. We used optogenetics to control visual cortex neurons with light. Different optogenetic stimulation protocols, using constant light, rhythmically modulated light or white-noise modulated light, all demonstrated that the investigated circuits predominantly resonate to stimulation in the gamma band. The observed gamma-band resonance renders visual cortical circuits most sensitive to gamma-rhythmic synaptic inputs. This in turn renders their spike output and the ensuing interareal synchronization gamma rhythmic.

This work was supported by DFG (SPP 1665, FOR 1847, FR2557/5-1-CORNET to P.F.; EXC 1086, DI 1908/5-1, DI 1908/6-1 to I.D.), BMBF (01GQ1301 to I.D.), EU (HEALTH-F2-2008-200728-BrainSynch, FP7-604102-HBP, FP7-600730-Magnetrodes to P.F.; ERC Starting Grant OptoMotorPath to I.D.), a European Young Investigator Award to P.F., the FENS-Kavli Network of Excellence to I.D., National Institutes of Health (1U54MH091657-WU-Minn-Consortium-HCP to P.F.), the LOEWE program (NeFF to P.F. and I.D.). Present address of I.D.: Optophysiology, Bernstein Center and BrainLinks-BrainTools, University of Freiburg, Albertstrase 23, 79104 Freiburg, Germany.

**Author contributions:** J.N, C.M.L., T.W., P.F. designed research; J.N, C.M.L., T.W., P.J., I.D., P.F. performed experiments; J.N., C.M.L., T.W. analyzed data; J.N., P.F. wrote the paper.

## Introduction

When visual cortex of an awake or lightly anesthetized subject is activated by appropriate stimuli, its neurons typically synchronize their activity in the gamma-frequency band, between 30 and 90 Hz (Gray et al., 1989; Kreiter and Singer, 1996; Hoogenboom et al., 2006). Very similar gamma-band synchronization has also been found outside visual cortex, e.g. in somatosensory and auditory cortex (Brosch et al., 2002; Bauer et al., 2006), motor cortex (Brown et al., 1998; Schoffelen et al., 2005; Ball et al., 2008), parietal and frontal cortex (Pesaran et al., 2002; Gregoriou et al., 2009; Lundqvist et al., 2016) and in hippocampus (Csicsvari et al., 2003; Colgin et al., 2009). Studies on gamma-band synchronization have investigated both its functional consequences and its mechanisms.

Studies on the functional role of visual cortical gamma-band synchronization have used in-vivo approaches to investigate relations to visual stimulation, task requirements and behavior. These studies suggest that visual gamma subserves, among other functions, object segmentation (Gray et al., 1989) and perceptual and attentional stimulus selection (Fries et al., 2002; Womelsdorf et al., 2006; Bosman et al., 2012; Grothe et al., 2012).

Experimental studies on the mechanisms underlying gamma-band synchronization have partly used in-vivo approaches. For example, intracellular recordings in anesthetized cat visual cortex revealed a type of cell, denoted “chattering cell”, that intrinsically generates gamma-rhythmic bursts when depolarized by current injection and that exhibits pronounced oscillations when visually stimulated (Gray and McCormick, 1996). Also, several in-vivo studies demonstrated that excitatory neurons lead inhibitory neurons during the gamma cycle by a few milliseconds (Csicsvari et al., 2003; Hasenstaub et al., 2005; Vinck et al., 2013). Yet, most experimental investigations of gamma mechanisms used in-vitro approaches, because they more readily allow intracellular recordings and current injections, as well as pharmacological manipulations (Whittington et al., 2000; Buzsáki and Wang, 2012; Salkoff et al., 2015).

Optogenetics, using Channelrhodopsin or related opsins, provides novel opportunities to investigate the mechanisms underlying the generation of rhythms, including the gamma rhythm. While a majority of optogenetic studies have exploited pathway or cell-type specific opsin expression to elucidate particular neural circuits, several studies capitalized on the excellent temporal resolution of optogenetic stimulation to investigate mechanisms underlying different neuronal rhythms. For example, the rhythmic optogenetic stimulation of mouse hippocampal or neocortical parvalbumin-positive (PV) interneurons leads to the indirect induction of theta resonance (Stark et al., 2013).

One seminal study used optogenetics in mouse somatosensory cortex to stimulate either PV interneurons or excitatory neurons (Cardin et al., 2009). When optogenetic pulse trains were given to PV interneurons, the network showed local field potential (LFP) resonance in a gamma-frequency band peaking around 40-50 Hz. By contrast, when the same pulse trains were delivered to excitatory neurons, resonance was strongest at 8 Hz and declined monotonically for increasing frequencies to vanish above 24 Hz.

We used optogenetics to investigate the resonance properties of visual cortex to provide further insights into mechanisms behind gamma-band synchronization among visual cortical neurons in vivo. We used visual cortex of the lightly anesthetized cat, a classical model system for research on vision and gamma-band synchronization. First, we tried three viral vectors and found that AAV5 fails to provide expression in the cat, whereas both AAV1 and AAV9 lead to robust expression. Constant optogenetic stimulation induced strong and sustained gamma-band activity. Rhythmic stimulation with pulse trains or sine waves at frequencies between 5 and 80 Hz revealed network resonance at 40 Hz or above. To investigate this resonance with greater spectral resolution, we applied optogenetic white noise stimulation, which confirmed resonance with a peak at 40-60 Hz.

## Materials and Methods

Eight adult domestic cats (*felis catus*; four females) were used in this study. All procedures complied with the German law for the protection of animals and were approved by the regional authority (Regierungspräsidium Darmstadt). After an initial surgery for the injection of viral vectors and a 4-6 week period for opsin expression, recordings were obtained during a terminal experiment under general anesthesia.

### Viral vector injection

For the injection surgery, anesthesia was induced by intramuscular injection of ketamine (10 mg/kg) and dexmedetomidine (0.02 mg/kg), cats were intubated, and anesthesia was maintained with N_2_O:O_2_ (60/40%), isoflurane (~1.5%) and remifentanil (0.3 µg/kg/min). Four cats were injected in area 17 and another four cats in area 21a. Rectangular craniotomies were made over the respective areas (Area 17: AP [0, -7.5] mm; ML: [0, 5] mm; area 21a: AP [0,-8] mm, ML [9, 15] mm). The areas were identified by the pattern of sulci and gyri, and the dura mater was removed over part of the respective areas. Three to four injection sites were chosen, avoiding blood vessels, with horizontal distances between injection sites of at least 1 mm. At each site, a Hamilton syringe (34 G needle size; World Precision Instruments) was inserted with the use of a micromanipulator and under visual inspection to a cortical depth of 1 mm below the pia mater. Subsequently, 2 µl of viral vector dispersion was injected at a rate of 150 nl/min. After each injection, the needle was left in place for 10 min before withdrawal, to avoid reflux. Upon completion of injections, the dura opening was covered with silicone foil and a thin layer of silicone gel, the trepanation was filled with dental acrylic, and the scalp was sutured.

In one cat, area 17 in the left hemisphere was injected with AAV1-CamKIIα-hChR2(H134R)-eYFP (titer 8.97*10^12^ GC/ml) and area 17 in the right hemisphere with AAV9-CamKIIα-ChR2-eYFP (titer 1.06*10^13^ GC/ml). In two cats, area 17 of the left hemisphere was injected with AAV1-CamKIIα-hChR2(H134R)-eYFP (titer: 1.22*10^13^ GC/ml). In one cat, area 17 of the left hemisphere was injected with AAV5-CamKIIα-ChR2-eYFP (titer 4*10^13^ GC/ml). In four cats, area 21a of the left hemisphere was injected with AAV9-CamKIIα-hChR2(H134R)-eYFP (titer: 1.06*10^13^ GC/ml). AAV1 and AAV9 viral vectors were obtained from Penn Vector Core (Perelman School of Medicine, University of Pennsylvania, USA), AAV5 viral vectors from UNC Vector Core (UNC School of Medicine, University of North Carolina, USA).

### Neurophysiological recordings

For the recording experiment, anesthesia was induced and initially maintained as during the injection surgery, only replacing intubation with tracheotomy and remifentanyl with sufentanil. After surgery, during recordings, isoflurane concentration was lowered to 0.6%-1.0%, eye lid closure reflex was tested to verify narcosis, and vecuronium (0.25mg/kg/h i.v.) was added for paralysis during recordings. Throughout surgery and recordings, Ringer solution plus 10% glucose was given (20 ml/h during surgery; 7 ml/h during recordings), and vital parameters were monitored (ECG, body temperature, expiratory gases).

Each recording experiment consisted of multiple sessions. For each session, we inserted either single or multiple tungsten microelectrodes (~1 MΩ at 1 kHz; FHC), or three to four 32-contact probes (100 µm inter-site spacing, ~1 MΩ at 1 kHz; NeuroNexus or ATLAS Neuroengineering) in area 21a and area 17. Standard electrophysiological techniques (Tucker Davis Technologies, TDT) were used to obtain multi-unit activity (MUA) and LFP recordings. For MUA recordings, the signals were filtered with a passband of 700 to 7000 Hz, and a threshold was set to retain the spike times of small clusters of units. For LFP recordings, the signals were filtered with a passband of 0.7 to 250 Hz and digitized at 1017.1 Hz.

### Photo-stimulation

Optogenetic stimulation was done with a 473 nm (blue) laser or with a 470 nm (blue) LED (Omicron Laserage). A 594 nm (yellow) laser was used as control. Laser light was delivered to cortex through a 100 µm or a 200 µm diameter multimode fiber (Thorlabs), LED light through a 2 mm diameter polymer optical fiber (Omicron Laserage). Fiber endings were placed just above the cortical surface, immediately next to the recording sites with a slight angle relative to the electrodes. Laser waveform generation used custom circuits in TDT, and timing control used Psychtoolbox-3, a toolbox in MATLAB (MathWorks) (Brainard, 1997).

For white-noise stimulation, the laser was driven by normally distributed white noise, with light intensities updated at a frequency of 1017.1 Hz. The total output intensity varied between sessions, with values in the range of 15-80 mW (46 recording sites in area 17 of 3 cats).

### Histology

After conclusion of recordings, approximately five days after the start of the terminal experiment and still under narcosis, the animal was euthanized with pentobarbital sodium and transcardially perfused with phosphate buffered saline (PBS) followed by 4% paraformaldehyde. The brain was removed, post-fixed in 4% paraformaldehyde and subsequently soaked in 10%, 20% and 30% sucrose-PBS solution, respectively, until the tissue sank. The cortex was sectioned in 50 µm thick slices, which were mounted on glass slides, coverslipped with an antifade mounting medium, and subsequently investigated with a confocal laser scanning microscope (CLSM, Nikon C2 90i, Nikon Instruments) for eYFP-labelled neurons.

#### Immunohistochemistry

In two cats, one with injections in area 17 and one with injections in area 21a, slices were processed as described above and additionally stained for parvalbumin (PV). To this end, slices were preincubated in 10% normal goat serum (Sigma Aldrich) with 1% bovine serum albumin (BSA) and 0.5% Triton X-100 in phosphate buffer (PB) for 1 h at room temperature to block unspecific binding sites. Slices were subsequently incubated in 3% normal goat serum with 1% BSA, 0.5% Triton X-100 and the primary antibody (rabbit anti-Parvalbumin, NB 120-11427, Novus Biologicals) over night at room temperature. After washing two times 15 min in PB, the slices were incubated with the secondary antibody (goat anti-rabbit Alexa Fluor 647, A-21244, Thermo Fisher Scientific) in 3% normal goat serum, 1% BSA and 0.5% Triton X-100 for 1 h at room temperature. Finally, slices were again washed in PB, coverslipped and imaged with a Zeiss CLSM, using a 25X water immersion objective.

### Data analysis

All data analysis was performed using custom code and the Fieldtrip toolbox (Oostenveld et al., 2011), both written in MATLAB (MathWorks).

#### Spike densities, spike autocorrelation histograms, LFP power spectra, and MUA-LFP PPCs

MUA rate was smoothed with a Gaussian (for constant light stimulation: SD =12.5 ms; for stimulation with pulse trains and sine waves: SD = 1.25 ms; in each case truncated at ± 2 SD) to obtain the spike density. To measure the MUA rhythmicity during optogenetic stimulation with pulse trains or sine waves of frequency f, the F1 component of the MUA was calculated as the amplitude of the Fourier transform at frequency f. To give F1 values of the different animals equal impact on the grand average, F1 values were z-scored within each animal by subtracting the mean and dividing by the SD of all F1 values of that animal.

The spike autocorrelation histogram (ACH) was calculated at 1 ms resolution with maximum time lag of 250 ms. Subsequently, the ACH was normalized by the triangle function tri(t) and the MUA rate, such that the ACH is expressed in units of coincidence/spike:

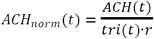

The triangle function is defined as

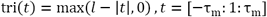

with *l* being the spike train length in seconds, and *r* being the mean spike rate in Hertz.

The ACH was smoothed with a Gaussian (SD = 0.5 ms, truncated at ±1.5 SD), and the F1 component of the ACH was calculated and normalized as for the MUA spike train.

LFP power spectra were calculated for data epochs that were adjusted for each frequency to have a length of 4 cycles and moved over the data in a sliding-window fashion in 1 ms steps. Each epoch was multiplied with a Hann taper, Fourier transformed, squared and divided by the window length to obtain power density per frequency. For the different stimulation frequencies f, LFP power is shown as ratio of power during stimulation versus pre-stimulation baseline (-0.5 s to -0.2 s relative to stimulation onset).

MUA-LFP locking was quantified by calculating the MUA-LFP PPC (pairwise phase consistency), a metric that is not biased by trial number, spike count or spike rate (Vinck et al., 2010). Spike and LFP recordings were always taken from different electrodes. For each spike, the surrounding LFP was Hann tapered and Fourier transformed. Per spike and frequency, this gave the MUA-LFP phase, which should be similar across spikes, if they are locked to the LFP. This phase similarity is quantified by the PPC as the average phase difference across all possible pairs of spikes. To analyze PPC as a function of frequency and time (Fig. 4 and 9), the LFP around each spike in a window of ±2 cycles per frequency was Hann tapered and Fourier transformed. PPC was then calculated for epochs of 100 ms length, i.e. using the phases of spikes in those epochs, moved over the data in a sliding-window fashion in 1 ms steps. To analyze PPC with higher spectral resolution (Fig. 5), the LFP around each spike in a window of ±0.5 s (Fig. 5F, lower frequencies) or ±0.25 s (Fig. 5F, higher frequencies) was Hann tapered and Fourier transformed to obtain the spike phase. For a given MUA channel, MUA-LFP PPC was calculated relative to all LFPs from different electrodes and then averaged.

#### Estimation of transfer function from recordings during white-noise stimulation

For each recording site and channel, the transfer function between the uniform white noise time series driving the laser and the simultaneously acquired local field potential was determined. The transfer function was estimated by Welch’s average periodogram method, separately per recording site and trial. It is the ratio of the cross spectral density between the input (laser) and the output (LFP) time series, and the power spectral density of the input (laser). To determine the white-noise driven resonance spectrum, the magnitude of the transfer function was computed for each recording site. The values from one such estimate demonstrate the transfer function for a single example recording site (Fig. 9D). In order to estimate the average transfer function across all recording sites, these magnitudes were normalized to equalize the total power. The normalized values across all recording sites were averaged to calculate the average spectrum (Fig. 9E).

#### Statistical testing

High-resolution spectra of LFP power changes and MUA-LFP PPC were compared between stimulation with blue light and control stimulation with yellow light (Fig. 5E,F). We calculated paired t-tests between spectra obtained with blue and yellow light, across recording sites. Statistical inference was not based directly on the t-tests (and therefore corresponding assumptions will not limit our inference), but the resulting t-values were merely used as a difference metric for the subsequent cluster-based non-parametric permutation test. For each of 10,000 permutations, we did the following: 1) We made a random decision per recording site to either exchange the spectrum obtained with blue light and the spectrum obtained with yellow light or not; 2) We performed the t-test; 3) Clusters of adjacent frequencies with significant t-values (p<0.05) were detected, and t-values were summed over all frequencies in the cluster to form the cluster-level test statistic. 4) The maximum and the minimum cluster-level statistic were placed into maximum and minimum randomization distributions, respectively. For the observed data, clusters were derived as for the randomized data. Observed clusters were considered significant if they fell below the 2.5^th^ percentile of the minimum randomization distribution or above the 97.5^th^ percentile of the maximum randomization distribution (Maris and Oostenveld, 2007). This corresponds to a two-sided test with correction for the multiple comparisons performed across frequencies (Nichols and Holmes, 2002).

## Results

### Transfection of cat visual cortex neurons by AAV1, AAV5 and AAV9

Recombinant adeno-associated virus (AAV) vectors are widely used as gene delivery tools (Vasileva and Jessberger, 2005). AAV-mediated expression of Channelrhodopsin-2 (ChR2) has been used in several mammalian species, including mice, rats and non-human primates (Diester et al., 2011; Gerits et al., 2015; Scheyltjens et al., 2015). In this study, three pseudo-typed AAVs (AAV1, AAV5 and AAV9) were tested in visual cortex of the domestic cat (*felis catus*). We injected AAVs carrying the gene for the expression of hChR2(H134R)-eYFP under the control of the Calcium/calmodulin-dependent protein kinase type II alpha chain (CamKIIα) promoter. Injections targeted either area 17, the cat homologue of primate area V1, or area 21a, the cat homologue of primate area V4 (Payne, 1993).

AAV5 was injected into area 17 of one cat and this did not result in visible ChR2-eYFP expression (Figure 1A). This failure of AAV5 expression is consistent with a previous AAV5 transduction study in cat cerebral cortex (Vite et al., 2003). By contrast, AAV1 and AAV9 injections into area 17, and AAV9 injections into area 21a resulted in robust ChR2-eYFP expression (Figure 1B-D). For both, AAV1 and AAV9, fluorescence showed a dependence on cortical depth, being strong in superficial layers, of medium strength in deep layers and relatively weak in middle layers (Figure 1B,C). Higher magnification revealed labeling of individual neurons (Figure 1B-D, right panels).

**Figure 1.**
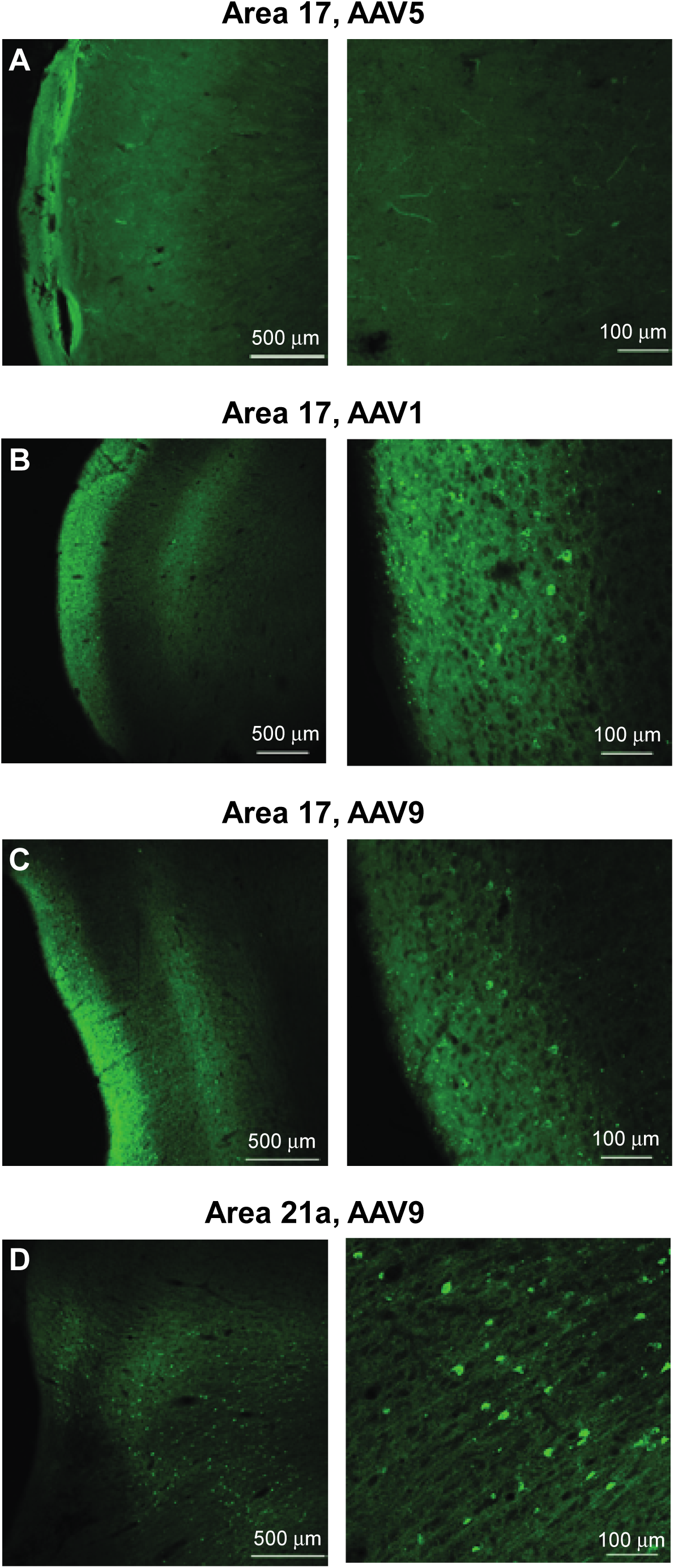
Transfection efficiency of different AAVs in cat visual cortex. Each pair of images shows the fluorescence at low resolution on the left side and at higher resolution on the right (see scale bars at lower right of each image). (A) Fluorescence in area 17 after injection with AAV5. Only auto-fluorescence is present, but no labeled cells. (B) Same as (A), but for area 17 injected with AAV1. (C) Same as (A), but for area 17 injected with AAV9. (D) Same as (A), but for area 21a injected with AAV9. Injections with AAV1 and AAV9 resulted in clear labeling of cells, indicative of successful ChR2 expression.

As described below, we find that optogenetic stimulation of the transfected neurons leads to network resonance in the gamma-band range. The generation of gamma-band activity has been linked to Parvalbumin-positive (PV+) interneurons. We therefore investigated, whether ChR2 was expressed in PV+ neurons. In two cats, we stained histological slices with fluorescence-marked antibodies against parvalbumin (Fig. 2A-F). One cat had been injected with AAV9-CamKIIα-ChR2-eYFP into area 21a, the other had been injected with AAV1-CamKIIα-hChR2(H134R)-eYFP into area 17. Across several slices and imaging windows of area 21a, we identified 182 unequivocally labeled neurons, which showed ChR2-eYFP expression or PV+ anti-body staining (Fig. 2A-C); of those, 73 were PV+ and 109 were ChR2-eYFP neurons, and there was zero overlap between these groups (Fig. 2G). Across several slices and imaging windows of area 17, we identified 282 unequivocally labeled neurons, which showed ChR2-eYFP expression or PV+ anti-body staining (Fig. 2D-F); of those, 154 were PV+ and 128 were ChR2-eYFP neurons, and again there was zero overlap between these groups (Fig. 2G).

**Figure 2.**
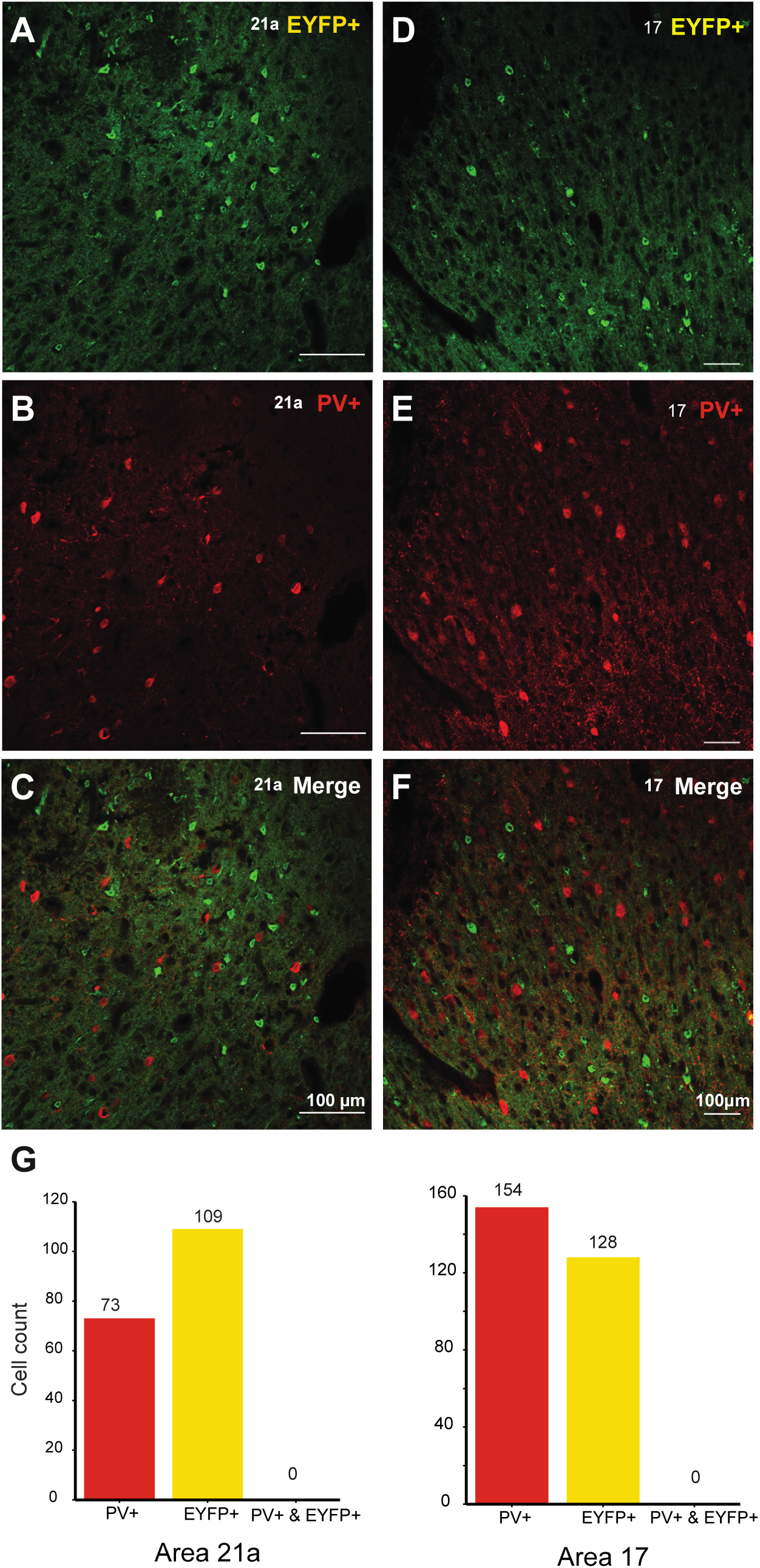
PV+ antibody staining. (A-C) Slice from area 21a. (D-F) Slice from area 17. (A, D) Cells expressing the fusion protein ChR2-eYFP. (B, E) PV+ antibody labeled cells. (C, F) Merged images, testing for neuronal co-labeling with ChR2-eYFP and PV+ antibody. No co-labeled neurons can be found. (G) Counts of PV+ labeled neurons, EYFP+ labeled neurons, and co-labeled neurons in area 21a (left side) and area 17 (right side).

### Neuronal responses to optogenetic stimulation after transfection with AAV1, AAV5 and AAV9

Between 4 and 6 weeks after virus injection, we performed terminal experiments under general anesthesia. The injected part of cortex was illuminated with blue light while neuronal spike and LFP activity was recorded from the optogenetically stimulated region. As mentioned above, one injection used AAV5 in area 17 of one cat and failed to show transfected neurons in the later histology. Correspondingly, the recordings in this case also failed to show any neuronal response to light application (Fig. 3A, B; pulses of 18 mW strength and 2 ms duration, applied in a regular 40 Hz pulse train). Firing rates following light pulses (in a window from 2 to 10 ms after light onset) did not differ significantly from rates immediately preceding the pulses (-10 to 0 ms) (Wilcoxon rank-sum test = 2503, *p* = 0.88, n = 5 sites). This was in stark contrast to responses in a cat injected with AAV1 and AAV9. In one cat, AAV1 was injected into area 17 in the left hemisphere (Fig. 3C,D), and AAV9 was injected into area 17 in the right hemisphere (Fig. 3E,F). Both injections led to strong optogenetic responses. Pulse trains of 20 Hz resulted in strong firing rate enhancements with a clear 20 Hz modulation. The peri-stimulus time histograms showed response latencies after light pulses of 3.9 ms (AAV1, Fig. 3D) and 3.6 ms (AAV9, Fig. 3F).

**Figure 3.**
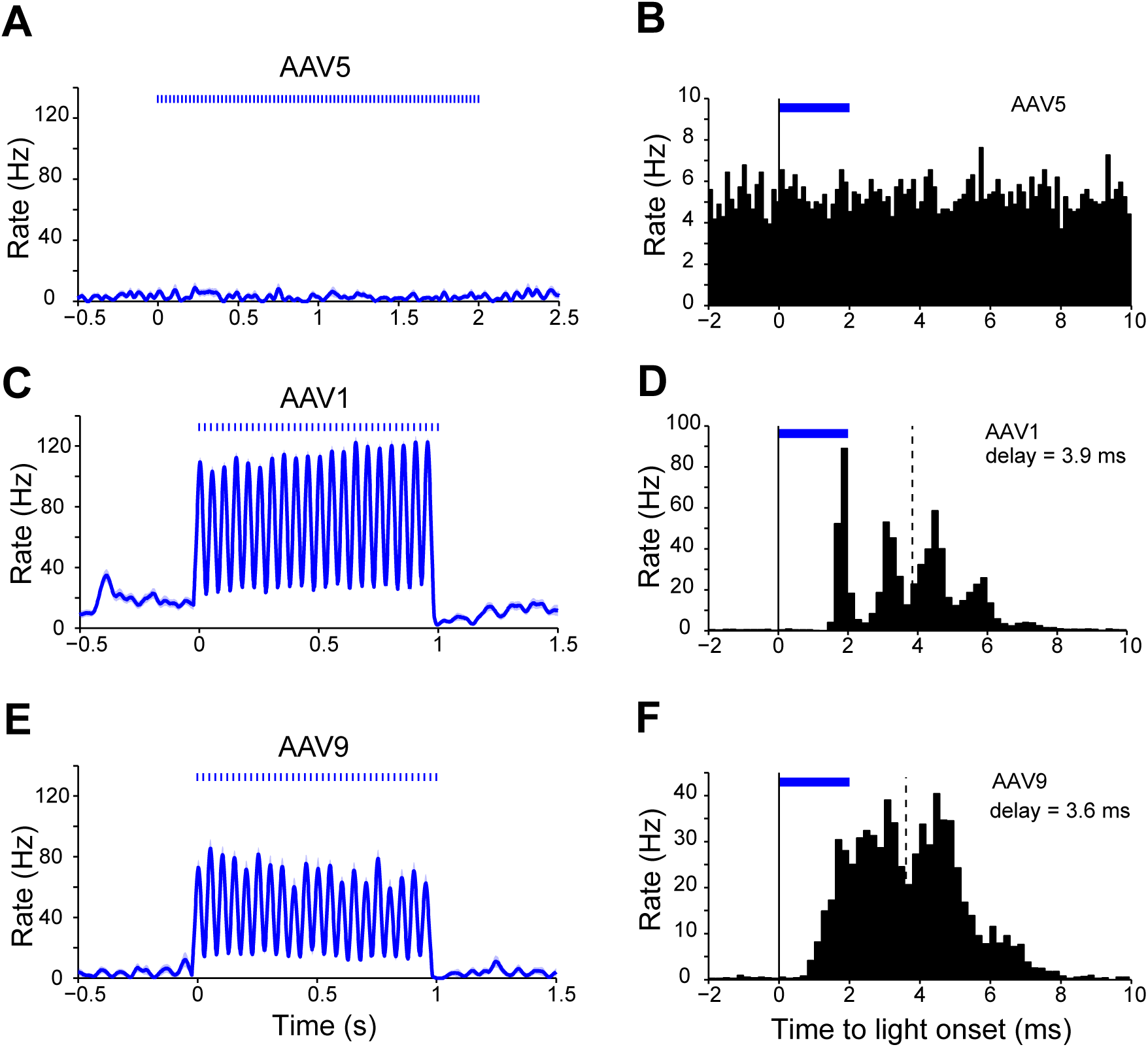
Neuronal responses to light stimulation after injection of AAV5, AAV1 or AAV9. (A) MUA in AAV5-injected area 17 during 40 Hz pulse-train light stimulation. Same MUA as shown in (A), averaged after aligning to the 2 ms light pulses. No optogenetic response was obtained after AAV5 injection. (C, E) MUA responses in area 17 after transfection with AAV1 (C) or AAV9 (E), and light stimulation with 20 Hz pulse trains. (D, F) Same data as (C, E), averaged after aligning to the 2 ms light pulses. MUA responses showed latencies of 3.9 ms and 3.6 ms (AAV1: 20 Hz, 10 -70 mW, N = 7 sites; AAV9: 20 Hz, 10 - 100 mW, N= 10 sites).

### Constant optogenetic stimulation induces neuronal gamma-band synchronization

Visual cortex shows particularly strong gamma-band activity in response to visual stimulation that is sustained and devoid of temporal structure (Kruse and Eckhorn, 1996). Thus, optogenetic stimulation of visual cortex might also be particularly suited to induce gamma if it is constant. Indeed, we have previously reported that constant optogenetic stimulation induces gamma-band activity in anesthetized cat visual cortex, in the context of an investigation of the gain-modulating effect of gamma (Ni et al., 2016). Here, we present a detailed analysis of gamma induced by constant optogenetic stimulation. Figure 4A shows an example LFP recording from area 21a of a cat transfected with AAV9, during one trial of optogenetic stimulation with 2 s of constant blue light. The raw LFP trace reveals strong optogenetically induced gamma, and the zoomed-in epoch illustrates that this emerged immediately after stimulation onset. Figure 4B shows the spike responses of this recording site for many interleaved trials of stimulation with blue or yellow light, confirming the selective optogenetic stimulation by blue light. Figure 4C shows the spike-triggered average of the LFP, demonstrating that spikes were locked to the LFP gamma-band component. The time-frequency analysis of both, LFP power (Fig. 4D, E) and MUA-LFP locking (Fig. 4F, G) showed a strong and sustained gamma-band peak for stimulation with blue light, that was absent for stimulation with yellow light.

**Figure 4.**
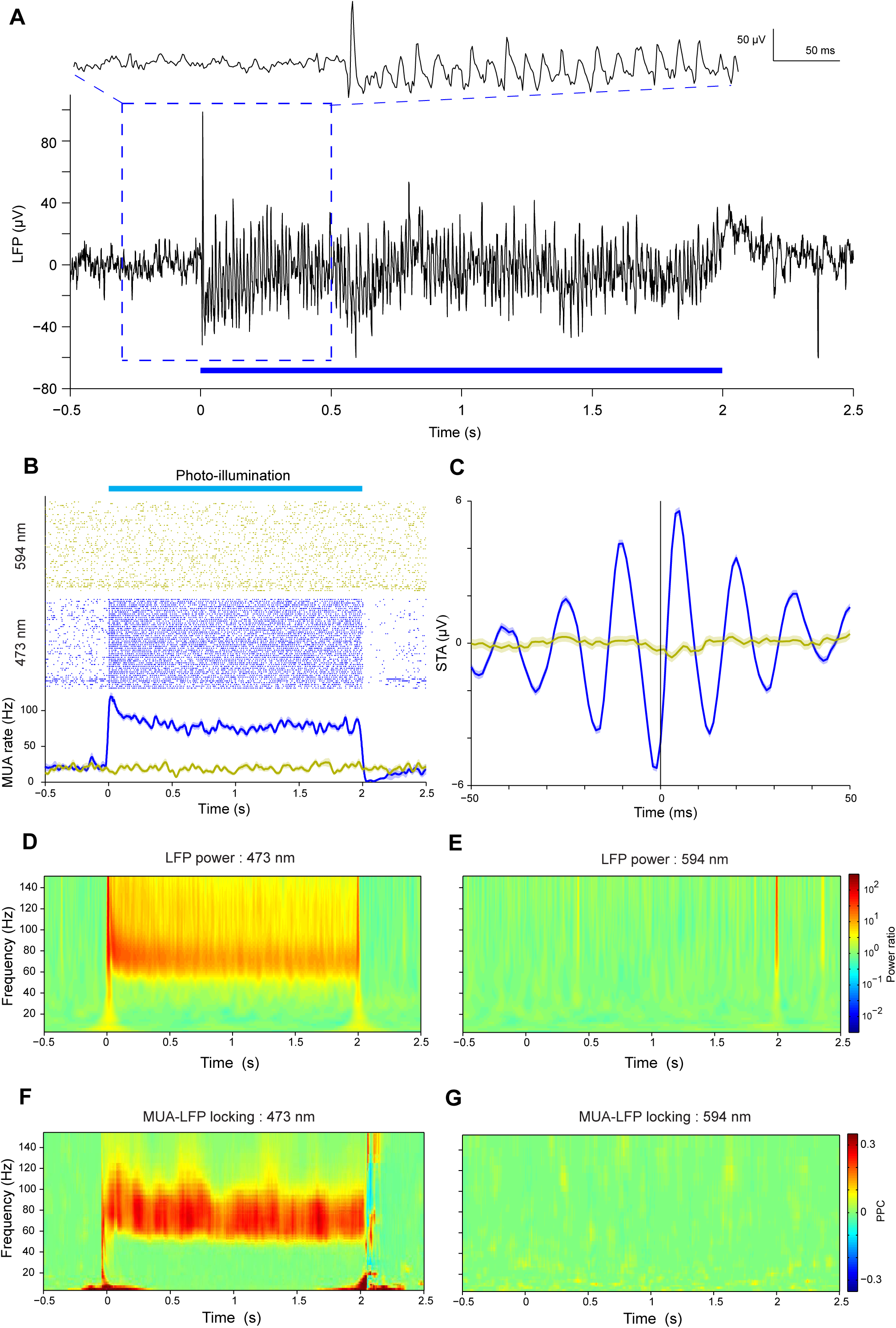
The effect of constant light stimulation on an example MUA and LFP recording site. (A) Constant-light induced gamma-band oscillation in the local field potential. (B) Constant-light induced MUA response. Blue: 473 nm wavelength light; Yellow: 594 nm wavelength light. The spike density was smoothed with a Gaussian function with σ = 12.5 ms, truncated at ±2σ. Shaded area indicates ±1 SEM. (C) Spike-triggered LFP. Shaded area indicates ±1 SEM. (D) TFR of constant-light induced gamma power, for 473 nm light. (E) Same as (D), but for 594 nm light as control. (F) TFR of constant-light induced MUA-LFP coherence. (G) The same as in (F), but for 594 nm light as control.

This pattern was found very reliably across all recording sites. Stimulation with two seconds of constant blue light, as compared to yellow control light, induced strong enhancements in firing rate, which were sustained for the duration of stimulation (Fig. 5A,D; Wilcoxon rank-sum test = 39581, *p<*0.0001, n =163 sites). The ratio of LFP power during stimulation versus pre-stimulation baseline showed an optogenetically induced gamma-band peak around 70 Hz (Fig. 5B,E; Wilcoxon rank-sum test = 14751, *p<*0.0001, n = 99 sites). We note that the gamma-band peak frequency varies across animals and recording sessions, as previously shown (Ni et al., 2016). The LFP gamma-power changes reflected changes in neuronal synchronization, because optogenetic stimulation also induced strong MUA-LFP locking in the gamma band, as quantified by the MUA-LFP PPC (Fig. 5C,F; Wilcoxon rank-sum test = 9389, *p<*0.0001, n = 84 sites). In addition to the induction of gamma-band activity, optogenetic stimulation also caused a power reduction between 6 and 12 Hz (Fig. 5E left panel for lower frequencies, note the scale is much smaller than for the higher frequencies shown in the right panel; Fig.5B inset). At the same time, it caused a reduction in MUA-LFP locking between 10 and 12 Hz (Fig. 5F and Fig.5C inset).

**Figure 5.**
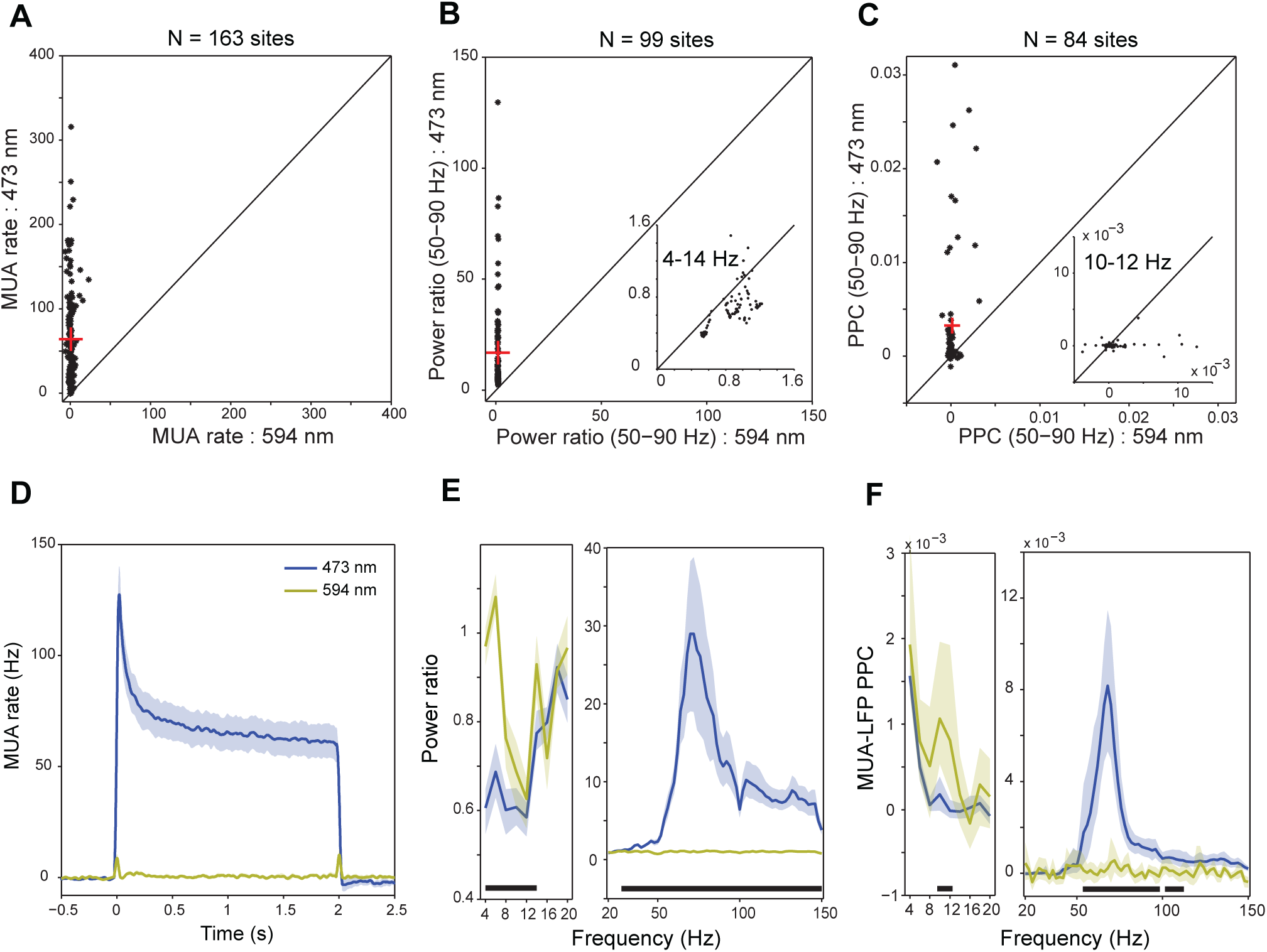
Group results for MUA rate, LFP power and MUA-LFP PPC induced by constant light. (A) Scatter plot of MUA rate (between 0.3 to 2 s) induced by constant light for both blue and yellow light condition. The red cross indicates the median value. (B) Scatter plot of power ratio (50-90 Hz) between blue and yellow light condition. The inset shows the same for 4-14 Hz, motivated by the analysis shown in (E). (C) Scatter plot of MUA-LFP PPC (50-90 Hz) between blue and yellow light condition. The inset shows the same for 10-12 Hz, motivated by the analysis shown in (F). (D) Averaged MUA spike density. Smoothed by a Gaussian function with σ = 12.5 ms and truncated for ±2σ. (E) Averaged power spectrum. (F) Averaged MUA-LFP PPC spectrum. (D-F) Blue (yellow) lines show data obtained with 473 nm (594 nm) light stimulation. Shaded areas indicate ±1 SEM, which is shown for illustration only. Black bars at the bottom indicate frequency ranges with statistically significant differences (*p <* 0.05), based on a cluster-level permutation test. (E, F) use ±0.5 s epochs for the analyses from 4 to 20 Hz, and ±0.25 s long epochs for the analyses from 20-150 Hz.

### Neuronal responses to optogenetic pulse train stimulation at different frequencies

To characterize the temporal response properties of the network to optogenetic stimulation of the transfected neurons, we applied pulse trains of different frequencies. Pulses always had a duration of 2 ms, and were repeated at frequencies of 5, 10, 20, 40, and 80 Hz. Pulse intensity was adjusted per recording site (see Materials and Methods) and was kept constant for a given site across the different pulse train frequencies. The analysis was limited to spike trains and excluded LFP, to avoid LFP artifacts caused by light stimulation. Pulse trains of all employed frequencies resulted in clear increases in firing rate, with strong rhythmicity at the pulse train frequency (Fig. 6). We calculated spike density functions, subtracted the baseline values and averaged them across recordings sites. Figure 6A-C shows those average spike densities for 10 Hz, 40 Hz and 80 Hz. We quantified their rhythmicity by calculating the Fourier transforms at the pulse train frequency (F1, see Materials and Methods). Figure 6D shows F1 for the different pulse train frequencies. Different stimulation frequencies led to different F1 components (one-way ANOVA, *p* = 2.3E-9, *F*_(4,200)_ = 12.9). F1 values showed a smooth dependence on pulse-train frequency, with a peak at 40 Hz. Auto-correlation histograms (ACHs) confirmed strong rhythmicity (Fig. 6E-G). F1 components of the ACHs also differed across frequencies (one-way ANOVA, *p* = 1.2E-6, *F*_(*4,200*)_ *=8.91*) and increased with frequency. Note that the ACH for 80 Hz stimulation suggested a 40 Hz subharmonic response.

**Figure 6.**
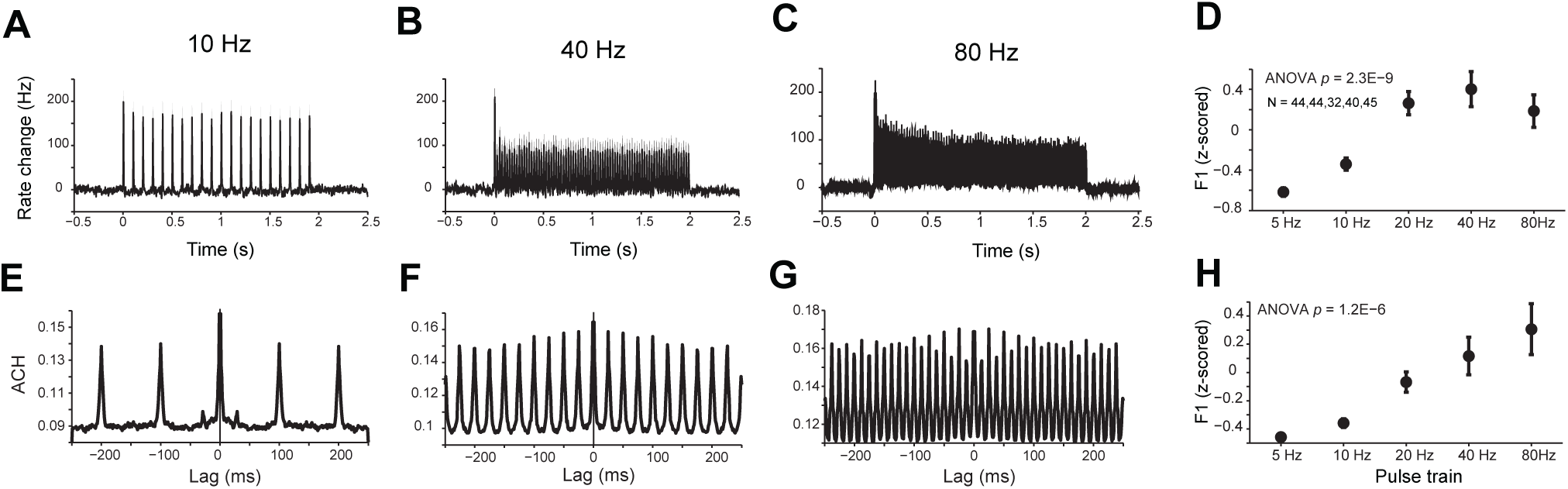
MUA resonance for optogenetic pulse-train stimulation. (A-C) MUA spike density (Gaussian smoothing with σ = 1.25 ms and truncated for ±2σ) for 10 Hz (A), 40 Hz (B) and 80 Hz (C) optogenetic pulse train stimulation, respectively. Data were averaged over all MUA recording sites. Error regions show ±1SEM, but are barely visible. (D) The F1 component of MUA spike density as a function of optogenetic pulse train frequency. (E-G) MUA auto-correlogram for 10 Hz (E), 40 Hz (F) and 80 Hz (G) optogenetic pulse train stimulation, respectively. Data were averaged over all MUA recording sites. (H) The F1 component of the MUA auto-correlogram (maxlag = 250 ms) as a function of the optogenetic pulse train frequency.

### Neuronal responses to optogenetic sine wave stimulation at different frequencies

The results so far suggest that the stimulated circuits resonate most strongly in the gamma-frequency band. Yet, for a fixed stimulation epoch, higher pulse train frequency imposed higher total light power onto the brain tissue. Therefore, we also employed optogenetic stimulation with sine waves of 5, 10, 20, 40 and 80 Hz, with total light power constant across frequencies. Results were similar to stimulation with pulse trains: The rhythmicity of the responses increased with stimulation frequency (Fig. 7A-H; one-way ANOVA: F1 of spike train, *p* = 3.6E-9, *F*_(4,295)_ = 12.17; F1 of ACH, *p* = 2.2E-14, *F*_(4,295)_ = 19.7). Note that the ACH for 80 Hz stimulation showed a substantial 40 Hz subharmonic response.

**Figure 7.**
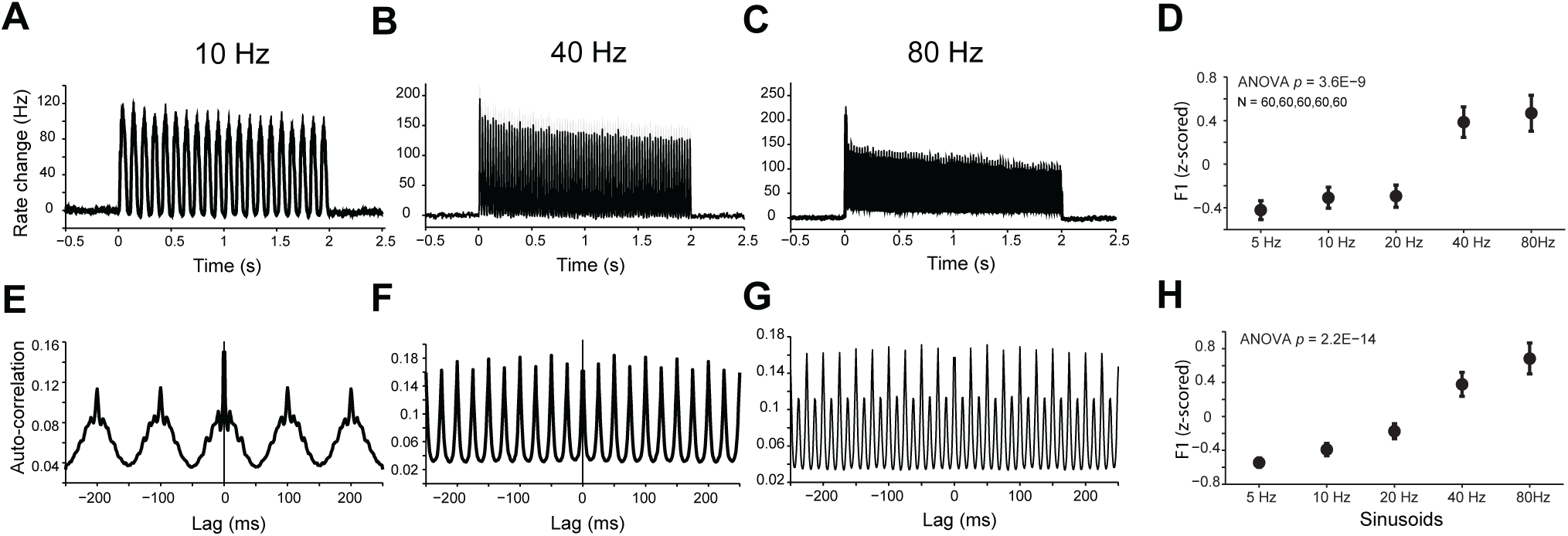
MUA resonance for optogenetic sine wave stimulation. (A-C) MUA spike density (Gaussian smoothing with σ = 1.25 ms and truncated for ±2σ) for 10 Hz (A), 40 Hz (B) and 80 Hz (C) optogenetic sine wave stimulation, respectively. Data were averaged over all MUA recording sites. Error regions show ±1SEM, but are barely visible. (D) The F1 component of MUA spike density as a function of optogenetic sine wave frequency. (E-G) MUA auto-correlogram for 10 Hz (E), 40 Hz (F) and 80 Hz (G) optogenetic sine wave stimulation, respectively. Data were averaged over all MUA recording sites. (H) The F1 component of the MUA auto-correlogram (maxlag = 250 ms) as a function of the optogenetic sine wave frequency.

### Neuronal response latencies to optogenetic stimulation with pulse trains and sine waves of variable frequencies

Next, we investigated the neuronal response latencies to optogenetic stimulation. This is highly relevant when optogenetic stimulation is used to produce temporal activation patterns at high frequencies. In addition, it provides a signature of true optogenetic stimulation. Optogenetic response latencies have typically been found on the order of 3-8 ms. Spikes detected at shorter latencies are suspicious of reflecting photo-electric artifacts (Cardin et al., 2010). To investigate response latencies, we averaged MUA responses aligned to the light pulses (Fig. 8A) or to the peaks of the sine wave light stimuli (Fig. 8B). Light pulses caused a small enhancement of MUA starting within approximately 1 ms after light onset and lasting a fraction of a millisecond, which most likely reflects light artifacts. The main response to the pulses followed later, starting at latencies after pulse onset of approximately 3 ms and peaking at 5.4-6.9 ms (Fig. 8C). During sine wave stimulation, the light was modulated between the respective maximal intensity and almost zero intensity. Thus, the light crosses the threshold for effective neuronal stimulation at an unknown intensity, and it is not possible to calculate response latencies in the same way as for the pulse trains. Therefore, we used a technique of latency estimation that has been developed in the study of synchronized oscillations, and that is based on the slope of the phase spectrum of the coherency between two signals (Schoffelen et al., 2005), in our case the light intensity and the MUA. Figure 8D shows this phase spectrum and reveals a strictly linear dependence of phase on frequency. The slope of this linear relationship allows to infer a latency of 5.5 ms, in close agreement to the values obtained for the different pulse train frequencies.

**Figure 8.**
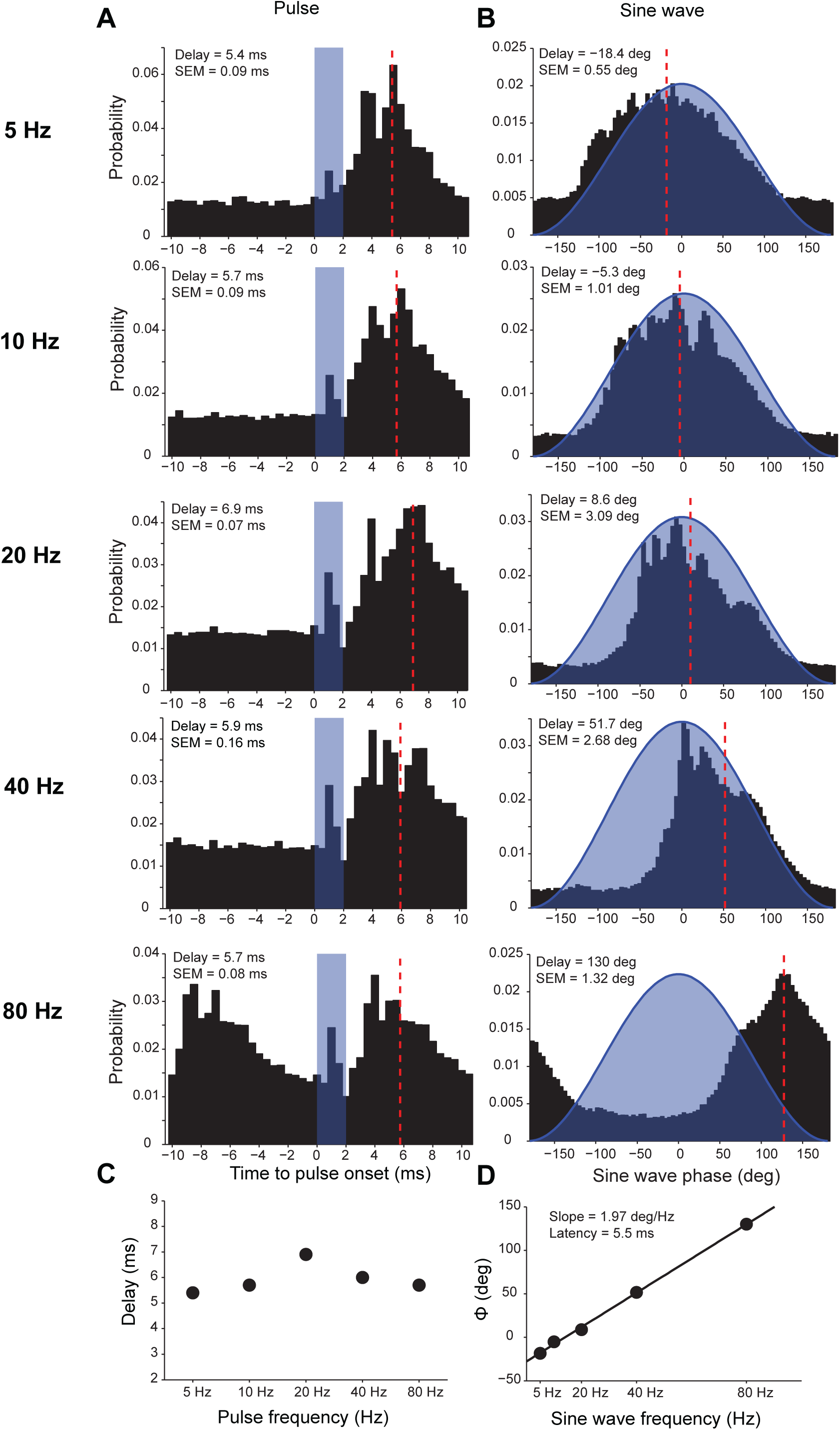
MUA response latencies to optogenetic pulse-train and sine-wave stimulation. (A) MUA spike probability, averaged over recording sites, as function of time relative to optogenetic stimulus pulse onset. The optogenetic pulse is indicated by the blue bar. Note the very short latency, very short-lived MUA enhancement around 1 ms, most likely reflecting light-induced artifacts in some of the recordings. (B) MUA spike probability, averaged over recording sites, as function of the phase of the optogenetic sine wave stimulation. The optogenetic sine wave is indicated by the blue-shaded region. (A, B) Data from the five pulse-train or sine-wave frequencies are shown in separate panels. MUA responses were fitted with Gaussians, and the resulting peak latencies are indicated by dashed red lines. Peak latencies and their SEM (estimated through a jackknife procedure) are also indicated as text insets. For sine-wave stimulation, latencies are expressed relative to the time of peak light intensity. (C) MUA peak latencies from (A) as a function of the pulse-train frequency. (D) MUA peak latencies from (B) as a function of the sine-wave frequency.

### Neuronal responses to optogenetic white-noise stimulation

The described neuronal responses to pulse trains and sine waves of different frequencies suggest that the network responds more strongly to rhythmic stimulation with higher frequencies, potentially with a peak in the gamma-frequency range. It would be interesting to characterize the spectrum of neuronal response to a large number of driving frequencies. Yet, testing neuronal responses to optogenetic stimulation at a sufficiently large number of individual frequencies to fully characterize the spectrum would require excessively long recordings. We therefore employed optogenetic stimulation with light intensities following a Gaussian random process (sampled at ≈1000 Hz) with a flat power spectrum (Fig 9). This white-noise stimulus contains the same energy at all frequencies up to 500 Hz. Recordings obtained during white-noise stimulation allow the estimation of the transfer function, which specifies for each frequency the strength of the neuronal circuit’s response given optogenetic stimulation at that frequency. Figure 9A shows an example LFP and MUA recording for an example trial of white-noise stimulation. The time-frequency analyses of the respective LFP power (Fig. 9B) and MUA-LFP locking (Fig. 9C), averaged over trials, showed sustained responses that peaked in the gamma-frequency range. The average transfer function between the white-noise stimuli and the example LFP recording site is shown in Figure 9D and reveals a dominant peak in the gamma band. Figure 9E shows the average transfer function across all recording sites stimulated with white noise (46 recording sites in area 17 of 3 cats), confirming a predominant peak in the gamma band.

**Figure 9.**
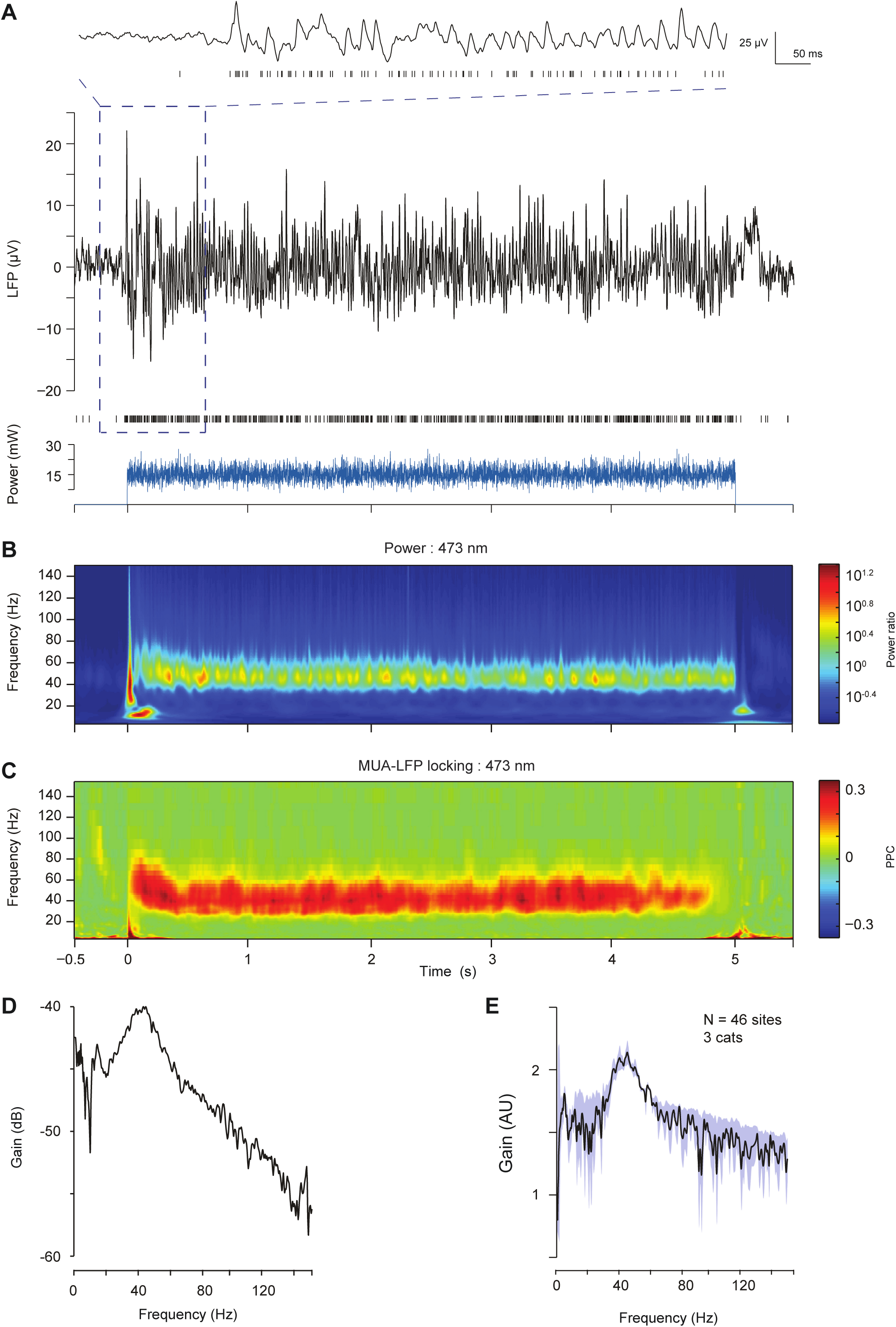
Cortical responses to white noise laser stimulation. (A-C) Example single trial LFP and MUA response to a white noise light stimulus. (A) The blue trace in the bottom panel shows the white-noise time course of laser intensity. The sequence of vertical lines above it indicates time points of MUA spike occurrence. The black continuous line shows the LFP. LFP and MUA for the epoch indicated by the dashed rectangle are shown at higher temporal resolution at the top. (B) LFP power as a function of time and frequency. (C) MUA-LFP PPC as a function of time and frequency. (D) The trial-averaged magnitude of the transfer function between white noise and the LFP shown in (A-C). (E) The magnitude of the transfer function, averaged over 46 LFP recording sites from 5 sessions in 3 cats. The blue region indicates ±1 SEM.

## Discussion

We investigate the response of anesthetized cat areas 17 and 21a to optogenetic stimulation with constant light, rhythmically modulated light and white-noise modulated light. Neurons were transfected to express Channelrhodopsin-2 (ChR2) through the injection of different adeno-associated viral vectors. We found that AAV5 fails to transfect cells in cat cortex. By contrast, both AAV1 and AAV9 resulted in robust and widespread transfection. ChR2 expression was under the control of a CamKIIα promotor, which is expressed in excitatory neurons. In agreement with that, immunohistochemical analysis in two cats found none of the transfected cells to express Parvalbumin, which suggests that optogenetic stimulation reached primarily or exclusively excitatory neurons. Optogenetic stimulation with constant light induced pronounced gamma-band activity, in agreement with several previous reports in other species and areas, as discussed in detail below; it also induced decreases in theta and alpha LFP power and alpha-band MUA-LFP locking. Rhythmic optogenetic stimulation with either pulse trains or with sinusoidal light modulation showed strongest resonance at 40 or 80 Hz. Stimulation with white-noise-modulated light revealed the full transfer function of the visual cortical circuit. This transfer function showed predominantly a peak in the gamma band, between 30 and 60 Hz.

The stimulation with light can cause artifacts in recordings with metal electrodes, as used here. We found artifacts in the LFP that were sizeable, yet constrained to the first few hundred milliseconds after light onset (data not shown). We also found artifacts in some of the MUA recordings, which were always constrained to the first 2 ms after light onset (Fig. 8A). The observation, that sustained optogenetic stimulation induces sustained gamma-band oscillations, is not due to artifacts, because gamma is sustained for the entire duration of optogenetic stimulation, long after the light-onset related artifact has ceased. The analysis of signals recorded with pulsed optical stimulation excluded LFPs, because under these conditions, our LFP recordings contained substantial artifact components, which were difficult to separate from neuronal components. By contrast, the analysis of MUA responses to pulsed light showed light-related artifacts that were small compared to the optogenetically driven neuronal responses (Fig. 8A). Finally, the analysis of signals recorded with white-noise optical stimulation most likely includes some artifacts in the LFP, yet these artifacts cannot explain the band-limited transfer function, because the power spectrum of light stimulation was by construction white.

Both, viral expression and light application was not homogeneous across cortical layers. Expression was stronger in superficial and deep layers than in middle layers, with a particular predominance in superficial layers. Optogenetic stimulation for all area 21a recordings and for most area 17 recordings was through surface illumination, leading to strongest light intensity in superficial layers. The combination of predominantly superficial expression with predominantly superficial light application likely led to a predominance of superficial neuronal activation. Superficial layers of macaque areas V1, V2 and V4, show substantially stronger visually induced gamma-band synchronization than infragranular layers (Buffalo et al., 2011; Xing et al., 2012). Thus, our finding of resonance in the gamma-frequency band might be partly due to a predominantly superficial localization of our optogenetic stimulation. Whether neuronal circuits in other layers resonate in different ways will require layer-specific expression and/or optical stimulation.

We found that visual cortical circuits resonate in the typical gamma-frequency band. Yet, across the different optogenetic stimulation protocols, we found variable peak frequencies. For constant optogenetic stimulation, the average gamma-band peak extended from 50-100 Hz. For rhythmically pulsed stimulation, only a limited set of frequencies was tested and strongest resonance was found at 40 or 80 Hz. For white-noise stimulation, the average transfer function peaked between 30 and 60 Hz. This variability in gamma peak frequency is likely due to a combination of factors. First, the gamma peak frequency is partly genetically determined and thereby varies across individuals (van Pelt et al., 2012). The different stimulation protocols analyzed here were applied to different subsets of cats, such that inter-individual differences in gamma peak frequency could lead to apparent differences between stimulation protocols. Second, the gamma peak frequency is likely affected by state changes, that can occur during anesthesia and might resemble changes in attention, which have been shown to modulate gamma peak frequency (Bosman et al., 2012). Third, the peak frequency of visually induced gamma-band activity is strongly affected by visual stimulus parameters with stimuli of higher salience typically leading to higher gamma peak frequency (Fries, 2015). Gamma peak frequency is e.g. reduced for stimuli that are of low contrast or masked by noise (Ray and Maunsell, 2010; Jia et al., 2013; Roberts et al., 2013). It is conceivable that similar effects occurred for the optogenetic stimulation employed in the present study. For example, rhythmically pulsed stimulation might be highly salient, leading to gamma resonance at the upper end of the gamma-frequency range; by contrast, white-noise stimulation might be of lower salience and similar to noisy visual stimulation, leading to lower gamma peak frequencies.

The finding that constant optogenetic stimulation of cortex induces sustained gamma-band activity is consistent with previous reports. Several in-vitro studies applied slowly ramping optogenetic stimulation to slices of somatosensory cortex or hippocampus and found that this induces strong and narrowband gamma-band activity (Adesnik and Scanziani, 2010; Akam et al., 2012; Crandall et al., 2015). Several in-vivo studies found that slowly ramping and/or constant optogenetic stimulation induces strong gamma-band activity that is less narrowband than in-vitro. When awake mouse frontal cortex receives sustained activation by means of a step-function opsin, this enhances power in a 50-90 Hz band and reduces power at 4-25 Hz (Yizhar et al., 2011). When awake macaque motor cortex is transfected to express the C1V1 opsin and stimulated with constant or slowly ramping light, it shows gamma-band activity of constant or varying peak frequency (Lu et al., 2015). When anesthetized cat visual cortex is transfected to express ChR2 and stimulated by constant light, it generates sustained gamma-band activity (Ni et al., 2016). This latter study used partly data from the same animals as used here, yet from different recording sessions. In the present study, we confirm that constant optogenetic stimulation of cat visual cortex induces gamma-band synchronization, and we add that it also reduces LFP power in the theta and alpha bands and MUA-LFP locking in the alpha band (Fig. 5E, F). These reductions in low-frequency LFP power and MUA-LFP locking are very similar to effects of visual stimulation and selective attention in awake macaque area V4 (Fries et al., 2008).

The findings with pulsed stimulation differ from earlier reports. A previous study used mouse Cre-lines to express ChR2 selectively in either PV-positive fast-spiking (FS) neurons or CamKIIα-positive regular-spiking (RS) cells (Cardin et al., 2009). Pulse-train stimulation of the PV-FS circuit led to LFP resonance in a 30-70 Hz frequency band. By contrast, pulse-train stimulation of the CamKIIα-RS circuit led to resonance for low frequencies, up to 24 Hz. The cells expressing ChR2 in the CamKIIα-Cre mice were 100% PV negative, consistent with exclusive expression in excitatory neurons. The present study in cat visual cortex cannot build on Cre lines to target expression to specific cell types. Any cell-type specificity of opsin expression is likely due to the employed CamKIIα promoter. Promoters generally control cell-type specific expression less tightly than Cre-driver lines. Nevertheless, the immuno-histochemical analysis in both area 17 and area 21a found 100% of the ChR2-expressing neurons to be PV negative. This strong exclusion of PV-positive neurons by the CamKIIα promotor might be specific for cat (visual) cortex. Previous studies in monkeys and rodents, using the same promoter, though sometimes with other AAV serotypes, found at least small proportions of labeled GABAergic cells (Nathanson et al., 2009b; Nathanson et al., 2009a; Galvan et al., 2016). Pulse-train stimulation applied to those putative excitatory cells in cat visual cortex revealed resonance in the gamma-frequency band, which was further corroborated by the white-noise stimulation. This is different from the low-pass resonance found with pulse-train stimulation of excitatory neurons in barrel cortex. This difference is likely due to the different modalities, somatosensation versus vision, and the different species, mouse versus cat, or a combination of both. In cat visual cortex, specific cells with the intrinsic and/or network-borne property of rhythmic bursting at the gamma rhythm have been described (Gray and McCormick, 1996; Cardin et al., 2005).

The current results suggest that input to (superficial) visual cortical circuits will be most effective if it is gamma rhythmic. If the input is spectrally broad, then resonance will be strongest to the gamma-band components. Local gamma-band synchronization is particularly strong in superficial layers (Buffalo et al., 2011; Xing et al., 2012), and inter-areal Granger-causal influences in the gamma-band are predominant along feedforward type projections, originating from superficial layers (Bastos et al., 2015a; Bastos et al., 2015b; Michalareas et al., 2016). Thus, local and interareal supragranular circuits might be tuned to render both, their output and their sensitivity to input, selective to the gamma band.

